# Rhythm of relationships in a social fish over the course of a full year in the wild

**DOI:** 10.1101/2022.03.11.483964

**Authors:** Ulf Aslak, Christopher T. Monk, Dirk Brockmann, Robert Arlinghaus

**Author notes:** These authors contributed equally to this work (Ulf Aslak, Christopher T. Monk). **Corresponding author:** UA, **Contact**. **Author Contributions:** UA, CTM, DB & RA conceived the ideas and developed the methodology, CTM and RA collected the data, UA and DB analysed the data, UA, CTM and RA led the writing and all authors contributed critically to the drafts and gave final approval to the manuscript.

## Abstract

Animals are expected to adjust their social behaviour to cope with challenges in their environment. Therefore, for fish populations, in temperate regions with seasonal and daily environmental oscillations, characteristic rhythms of social relationships should be pronounced. To date, most research concerning fish social networks and biorhythms has occurred in artificial laboratory environments or over confined temporal scales of days to weeks. By contrast, little is known about the social networks of wild, freely roaming fish, including how seasonal and diurnal rhythms modulate social networks over the course of a full year. The advent of high-resolution acoustic telemetry enables us to quantify detailed social interactions in the wild over time-scales sufficient to examine seasonal rhythms at whole-ecosystems scales. Our objective was to explore the rhythms of social interactions in a social fish population at various time-scales over one full year in the wild by examining high-resolution snapshots of dynamic social network. To that end, we tracked the behaviour of 36 adult common carp, *Cyprinus carpio*, in a 25 ha lake and constructed temporal social networks among individuals across various time-scales, where social interactions were defined by proximity. We compared the network structure to a temporally shuffled null model to examine the importance of social attraction, and checked for persistent characteristic groups (“friendships”) over time. The clustering within the carp social network tended to be more pronounced during daytime than nighttime throughout the year. Social attraction, particularly during daytime, was a key driver for interactions. Shoaling behavior substantially increased during daytime in the wintertime, whereas in summer carp interacted less frequently, but the interaction duration increased. Characteristic groups were more common in the summer months and during nighttime, where the social memory of carp lasted up to two weeks. We conclude that social relationships of carp change diurnally and seasonally. These patterns were likely driven by predator avoidance, seasonal shifts in lake temperature, visibility, forage availability and the presence of anoxic zones. The techniques we employed can be applied generally to high-resolution biotelemetry data to reveal social structures across other fish species at ecologically realistic scales.

## Introduction

Animals are faced with continual exogenous oscillations of their environment and endogenous oscillations of their own physiology, over various timescales (Hastings, 2010). For example, all animals must cope with daily oscillations driven by the Earth’s rotation around its axis, ~30 day oscillations driven by the lunar cycle and yearly oscillations driven by the Earth’s elliptical orbit around the sun. Animals must also cope with internal oscillations driven by, for example, a heartbeat at very short time scales, or a reproductive cycle at longer, maybe even seasonal time scales. Furthermore, to feed and survive animals must also track the responses of their predators and prey to environmental oscillations (Vandermeer, 2004). Importantly, in ectotherms such as fish, ecosystem metabolism (e.g., productivity of resources and thermal environment) in the temperate zone reacts strongly to seasonal and daily changes in light and temperature, which causes periodic variation in the availability of food and the distribution of habitats (Hunt, Jardine, Hamilton, & Bunn, 2012; Stæhr & Jensen, 2007). Accordingly, animals must constantly respond to oscillations in their biotic environment throughout the food web as resource requirements and availability, danger and shelter, and reproductive opportunities oscillate across various frequencies to find food and shelter and avoid mortality.

Animals have adopted a number of physiological and behavioural strategies to cope with periodic fluctuations of the environment over time. The circadian clock, for example, is an inner oscillator synchronized with solar time that appears universally across taxa (Bernard, Gonze, Čajavec, Herzel, & Kramer, 2007; Dunlap, 1999; Edgar et al., 2012); it influences metabolism (Kohsaka & Bass, 2007), hormones (Leatherland & McKeown, 1973) and ultimately behavior (Naylor, 1988). To survive resource fluctuations over yearly time-scales organisms may employ strategies like hibernation or other forms of metabolic depression (Ruf & Geiser, 2015). For most animals, and for ectotherms in particular, temperature is a critical resource they must adapt to (Magnuson, Crowder, & Medvick, 1979). In fishes, all biological processes are influenced by exogenously triggered temperature, including enzyme activity, metabolism, digestion, and feeding rate, leading to a strict dependency on warm waters to grow and reproduce (Conover & Present, 2016; Shultz, Reynolds, & Conover, 1996). In addition to modifying their physiology, animals also have the option to alter their behaviour in response to environmental changes (Holland, Brill, Chang, Sibert, & Fournier, 1992), which may take the form of migrating to more favourable habitats (Somveille, Rodrigues, & Manica, 2015), and importantly, animals may also change their response to other conspecifics by becoming more or less social (Monk et al., 2018).

There are costs and benefits to both pro-social and anti-social behaviour, which depend on an organism’s environment and resource requirements (Monk et al., 2018; Snijders, Kurvers, Krause, Ramnarine, & Krause, 2018; Wiens, 1976). Group living can allow for increased predator avoidance (Foster & Treherne, 1981; Landeau & Terborgh, 1986; Pulliam, 1973), faster ability to find rare or mobile resources (Hills et al., 2015; Magurran & Higham, 1988; Pitcher, Magurran, & Winfield, 1982), increased ability to hunt large prey, and better conservation of resources (Gilbert, Robertson, Le Maho, Naito, & Ancel, 2006). However, living in groups also comes at the cost of sharing resources among conspecifics (Bertram, 1978) or increased transmission of parasites and pathogens (Côté, Poulin, & Zealand, 1985). Under certain conditions it is therefore, better to be solitary and defend a territory (J. Brown, 1968; Bryant & Grant, 1995) or behave nomadically (Eklöv, 1992). Hence, as the abiotic and biotic environments as well as the internal physiological state of an animal oscillate we expect to observe periodic patterns in social behaviour across various time-scales. The behavioural reactions that occur most likely depend on the evolutionary adaptions of particular species, modified by local environmental conditions. For example, killer whales, *Orcinus orca*, increase their sociality with increasing resource abundance (Franks et al., 2012), while chacma baboons, *Papio hamadryas ursinus*, increase their sociality when resources become scarce (Henzi, Lusseau, Weingrill, Van Schaik, & Barrett, 2009). The environment-species interaction also dictates among population differences in social responses to the environment within the same species. For example banded killifish, *Fundulus diaphanus*, form larger groups when a predator is detected, but reduce their group size when food is available (Hoare, Couzin, Godin, & Krause, 2004).

To date, there have been few explorations into the temporal dynamics of animal social networks (Blonder, Wey, Dornhaus, James, & Sih, 2012; Pinter-Wollman et al., 2014), and none have been able to explore the dynamics across the full spectra of timescales expected within a full growing season in the temperate zone. The lack of long-term (i.e., multiple months), high-resolution animal social network data in the wild is a result of the immense challenge in collecting it at sufficient spatial-temporal resolutions (Krause et al., 2013). Most long-term datasets are generated through consistent periodic visual observations of interactions among identifiable animals (Henzi et al., 2009; Wittemyer, Douglas-Hamilton, & Getz, 2005). There is a particular lack of long-term social network data for fish because making long-term underwater observations has not been possible until recently. Initial work in the wild has however shown that certain fish species tend to be detected in groups of characteristic individuals (Hay & McKinnell, 2002; Klimley & Holloway, 1999; Ward et al., 2002; Wilson et al., 2014), but this pattern is not universal across species (Helfman, 1984). The lack of long term studies of fish in the wild is problematic because fish have been used widely as valuable model organisms to study social behaviours, such as shoaling (Wilson, Croft, & Krause, 2014), but most research in social behavior among fishes has occurred over short time periods of a few weeks, and often in non-naturalistic laboratory environments (Wilson et al., 2014). Evidence from non-human animal studies shows that social behavior is usually more dependent on ecology than taxonomy (Lefebvre, Palameta, & Hatch, 1996), raising doubts as to whether our lab-scale understanding is transferable to the wild (Sutter & Arlinghaus, et. al., 2012; Niemelä & Dingemanse, 2014). Consequently, very little is known with certainty about the relationships and social lives of wild-living fishes in large populations over long time-scales of multiple months in the wild.

Most species of fish must engage in social relationships for at least a portion of their life, for example during mating and shoaling for predator avoidance (Pitcher, 1986; Shaw, 1978). To increase fitness, many species socialize to establish hierarchies, exchange information and avoid predators (Seppälä, Karvonen, & Valtonen, 2008; Suboski & Templeton, 1989). Research in social learning has demonstrated a number of advanced social behaviors. For example, it has been shown that fish can learn escape routes from each other (C. Brown & Laland, 2002), infer social hierarchies by observing fights (Grosenick, Clement, & Fernald, 2007), recognize individual conspecifics (Griffiths & Magurran, 1997), and when given conflicting social cues tend to value public information over immediate social information (Coolen, Ward, Hart, & Laland, 2005). More complex social behaviors such as cooperation, establishment of partnerships and reciprocation of behaviors that require risk taking (Croft et al., 2006; Granroth-Wilding & Magurran, 2013; Milinski, Kulling, Kettler, & Bern, 1990), indicate that fish have well developed social cognition capabilities (Bshary, Gingins, & Vail, 2014).

Today, with advanced computational methods and modern tracking technology, such as high-resolution, precise, acoustic telemetry it is possible to collect snapshots of social interactions at several second frequencies over years in the wild (Nathan et al., 2022; Baktoft, Zajicek, Klefoth, Svendsen, & Jacobsen, 2015; Guzzo et al., 2018; Krause et al., 2013), both underwater (Lennox et al., 2017) and in terrestrial environments (Wilmers et al., 2015). A suite of techniques are now available for constructing and analyzing social networks generated from acoustic data (Blonder et al., 2012; Finn et al., 2010; Jacoby & Freeman, 2016; Mourier, Brown, & Planes, 2017). With a full-year dataset containing high-resolution mobility traces of 36 common carp (*Cyprinus carpio*) in a small lake, we for the first time explore the “rhythm of relationships” in nature at a whole ecosystem scale for a fish. We ask a basic question: how do seasons and daytime modulate the social behavior of a fish species described as highly social from laboratory contexts (Huntingford et al. 2010)?

Wild common carp are long-lived, omnivorous, typically benthivorous, warmwater cyprinids, native to eastern Europe and Asia. The species has been domesticated for aquaculture purposes as early as 2000 years ago and globally introduced into the wild where it forms feral populations (Balon, 2004). Carp have permanently established widely across the freshwater ecosystems around the globe (Howes, 1991; Parameswaran, Alikunhi, & Sukumaran, 1972; Vilizzi, 2012). They constitute a key fisheries resource for both commercial (FAO, 2018) and recreational fisheries (Arlinghaus & Mehner, 2003). Yet, appreciation of carp is not global. The species is also considered a pest in certain regions where it is originally non-native, such as North America (Bajer, Sullivan, & Sorensen, 2009), Australia (Taylor, Tracey, Hartmann, & Patil, 2012) and parts of Europe, such as Spain (Benito, Benejam, Zamora, & García-Berthou, 2015) as they can disturb aquatic macrophytes, leading to loss of water clarity and increased nutrient concentrations (Bajer et al., 2016). Accordingly, learning about the social behaviour of carp could help to improve both carp fisheries management (Klefoth, Skov, Kuparinen, & Arlinghaus, 2017), and eradication techniques (Bajer, Chizinski, & Sorensen, 2011).

The behaviour of carp is variable, both among populations (Hennen & Weber, 2014; Benito et al., 2015; Weber, Brown & Willis, 2016) and among individuals (Monk & Arlinghaus, 2017; Pollux, 2017). Increasing levels of domestication in stocked fish are known to increase the carp’s boldness (Klefoth, Skov, Krause, & Arlinghaus, 2012) and foraging activity and ingestion rates (Klefoth, Pieterek, & Arlinghaus, 2013). Carp are also known to be a social species, frequently found in groups (Bajer et al., 2011; Johnsen & Hasler, 1977; Osborne, Ling, Hicks, & Tempero, 2009), and can learn by social facilitation (Zion, Barki, Grinshpon, Rosenfeld, & Karplus, 2007). Carp are generally thought to occupy littoral habitats during the spring and summer and move to deeper waters to overwinter in larger groups (Armstrong et al., 2016; Johnsen & Hasler, 1977; Jones & Stuart, 2009; Penne & Pierce, 2008). Carp are also known to show marked diurnal behavioural patterns in the spring and summer months. For example, non-native carp that became established in a reservoir in the Ebro catchment in Spain were relatively inactive in deep hypoxic waters at nighttime, possibly refuging from nocturnal predators (in particular catfish, *Silurus glanis*), but became more active in shallow waters during the daytime (Benito et al., 2015). By contrast, in other populations, carp have been observed to increase food consumption in the nighttime (Bajer, Lim, Travaline, Miller, & Sorensen, 2010; Proske, 1972), strongly indicating that diel patterns of behavior will vary with local ecological conditions.

Our objective was to explore the social relationships of a population of carp over one full year at a whole ecosystem scale, and to identify the relevant time-scales of carp social behavior in the wild under realistic ecological scales. To that end, we recorded three-dimensional positions of carp in a whole-lake using high resolution acoustic telemetry, and inferred the temporal social network of carp by logging proximity events. We describe how seasons and daytime influence both mobility and social behavior, perform statistical tests to assess the degree to which time spent together is explained by social attraction given that other ecological factors (e.g., local food availability) may drive co-location of two individuals, and finally study the social network at varying time-scales to measure the persistence of community structure over time. Having found that there is significant local clustering in the social network, we investigate whether these groups, or communities, that emerge on short time-scales and give rise to local clustering, persist over time. By analyzing the persistence of clustering over time we aim to get at the more fundamental question which is whether there is social memory in the system that drives the groups of fish to meet repeatedly.

## Methods

### Study site

Kleiner Döllnsee (52°59’41.9”N, 13°34’56.4”E) is a 25 ha lake in northern Brandenburg, Germany, classified as eutrophic with a total phosphorous concentration of 38 *μg* L^-1^ at spring overturn. The average depth during the study period was 4.4 m, while the maximum depth is 7.8 m and the secchi depth was 1.97 ± 0.61 m (mean ± standard deviation). Reeds (*Phragmites australis*) form a belt of growth around the lake. Between May and October the lake stratifies turning layers below ca. 4 m anoxic (See figures SI1 and SI2). Kleiner Döllnsee hosts 14 fish species typical for mesotrophic to slightly eutrophic natural lakes in German lowlands (Eckmann, 1995). Top predators include introduced European catfish (*Silurus glanis*), and native northern pike (*Esox lucius*) and Eurasian perch (*Perca fluviatilis*). Carp competitors include an abundant populations of large common bream (*Abramis brama*), tench (*Tinca tinca*), rudd (*Scardinius erythrophthalmus*), roach (*Rutilus rutilus*) and white bream (*Blicca bjoerkna*). Through recent nutrient increases, the submerged macrophyte coverage has been declining in the lake and is now restricted to near shore locations and macrophytes taller than 10 cm covered 9.2% of the lake area.

### Telemetry system

The lake has been equipped with 20 submerged (~2 m) high-resolution acoustic telemetry receivers (WHS 3050; 200 kHz; Lotek Wireless Inc., Newmarket, Ontario, Canada) distributed at fixed locations throughout the lake (described in detail in Baktoft et al. 2015). The system allows whole-lake positional telemetry in 3-D at high spatio-temporal resolution with several position fixes per minute depending on transmitter burst rates (Baktoft et al. 2015). Location datapoints are estimated by hypertriangulation of ultrasonic signals originating from surgically implanted transmitters. Median location accuracy throughout the lake is 3.1 m (Baktoft et al., 2015). Macrophytes, known to strongly attenuate acoustic signals, were scarce during the study period; thus we can reason that most signal loss occurred when the fish swam among the reed close to shore. However, significant decreases in data yield did occur over some periods in warmer summer months, indicating improved telemetry performance during the cooler periods of the year (Fig. 2A). Average across the year data yield was about 40%. For a full description of the system and its performance see Baktoft et al., (2015).

### Carp population

All carp recorded in the dataset were hatchery born and bred in earthen ponds as is typical in many European fisheries where carp are stocked after being raised in pond aquaculture. In June 2014, 91 carp with transmitters implanted (0.3% to 2.2% body mass) were released to Kleiner Döllnsee. Due to tag loss, known to be a prevalent problem in carp tagging (Daniel, Hicks, Ling, & David, 2009; Økland, Hay, Naesje, Nickandor, & Thorstad, 2003), an additional 24 tagged carp were released in September 2014. Of these 115 carp known to have been introduced to the lake, between 25-36 (540 ± 79 cm total length, mean ± standard deviation; see Fig SI1 for individual level data) were successfully monitored throughout all of 2015 for an entire year (Fig. 1). The rest experienced transmitter loss. Tagging-induced mortality was extremely low as revealed by recaptures that had lost tags but were alive.

**FIG. 1.**
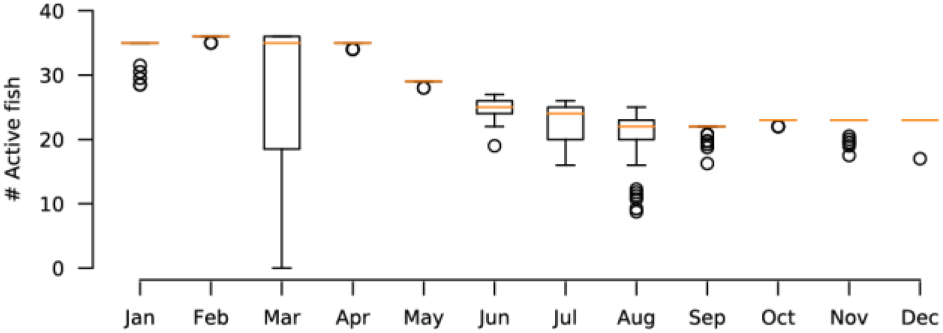
Distribution of number of fish with successfully recorded mobility on each day, across the year. At the beginning of the year 36 fish are active and at the end of the year 23 are active.

For transmitter implantation, carp were anaesthetized using a 9:1 EtOH:clove oil solution added at 1mL L^-1^ (Carl Roth, Karlsruhe, Germany). All surgical tools and acoustic telemetry tags were sterilized with a mixture of tap-water and 7.5% povidone-iodine (PVP; Braunol®; B. Braun, Kronberg, Germany) before each transmitter implantation. We implanted the transmitters (model MM-M-TP-16-50, dimension: 16 by 85 mm, wet weight: 21 g; Lotek Wireless, Newmarket, Canada) into the body cavity (see (Klefoth, Kobler, & Arlinghaus, 2008) for procedures), and each fish received 4-5 sutures using PDS-II adsorbable monofilament suture material and FS-1 3-0 needles (Ethicon, USA). Following recovery from surgery the fish were immediately released into the study lake. The burst frequency of the transmitters was five seconds, and the transmitters were equipped with a temperature sensor, recording once per minute, and a pressure sensor to record depth at all other transmissions.

### Inferring social networks

Because we did not directly measure social interactions between fish, we had to infer it using mobility traces. In the current analysis, we used persistent proximity as a proxy for contact. We employed several post-processing techniques to increase accuracy and data yield to produce the best reconstruction of the temporal contact network as possible. First, we resampled the location data from 5 s to 15 s to remove noise and recover potentially lost measurements by applying a 30 s median filter in 15 s increments across the location trajectory of each fish. We then measured pairwise distance in each time-bin, resulting in 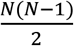 time series of inter-fish distance. Each series would have some missing values which we linearly imputed by up to 30 minutes. To get the times at which links were on and off we thresholded the distance series at 10 m. Finally, we applied a filter which removed singleton contact events and clustered consecutive ones with short breaks (up to 5 minutes). Figure 5 shows two examples of distance time series, how we threshold them and how that results in a link activity time series.

Based on individual temperature and location measurements of the *N* = 36 carp we computed, for each fish, *i*, on each day the local temperature, *T_i_*, the distance from shore, *d_i_*, the Shannon entropy, *S_i_* = ∑*_m_C_m_* log *c_m_*, where *c_m_* is the fraction of time spent in lake area *m* (when the lake is split into 10 m x 10 m cells), the velocity, *v_i_*, and the depth, *h_i_*. From the inferred social network we measured for each pair, *n*, the average interaction duration, *τ_n_*^+^, the average time between interactions, *τ_n_^−^*, and the interaction probability, *p_n_*. Quantities were estimated separately for daytime and nighttime once per solar cycle and reported as population averages, denoted by dropping the node/link index.

We, furthermore, measured the local clustering coefficient as a population average for each monthly aggregated social network split into day and night. The average local clustering in a network is bounded between zero and one, and reflects the tendency for triangles, or triads, to form in the network (Saramäki, Kivelä, Onnela, Kaski, & Kertesz, 2007). Intuitively, if a node has a high local clustering coefficient its neighbors are highly interconnected. Triads are indicative of community structure and informs about social behavior at the group-scale (Wasserman & Faust, 1994; Scott, 2000).

### Null model of social attraction

Since our inferred social network builds on the assumption that co-location equates interaction, it is natural to wonder how much of that interaction is due to “social attraction” – meaning very broadly that fish go to specific places because there are other fish there – and how much is due to the environment driving the fishes to visit the same places at the same times. Indeed, it is plausible that the population is entirely non-social and any co-location is due to similar resource use (e.g., habitat choice). To assess the impact of social attraction on interaction, we created a shuffled dataset using a null model that time-shifts the mobility trace of each fish independently by a random number of whole days between zero and six. In the shuffled dataset, any potential correlation in the location traces of two individuals due to their social attraction was broken (Spiegel, Leu, Sih, & Bull, 2016). The data with individually shifted mobility tracks then modeled a mobility pattern where each pair swims entirely independent of each other, effectively breaking location dependencies that may have existed due to social attraction. Hence, we got an estimate of the background level of that statistic due to habitual space use. This was, furthermore, a very strong null model since some fraction of fish pairs would be shifted the same amount (statistically: 7 (1/7)^2^ = 14.3%), thus not removing all inter-pair dependency. Effect sizes would therefore be slightly underestimated. We then measured and compared the raw number of interactions as well as the average local clustering coefficient in the real and shuffled data. We used this null model because, intuitively, if carp were truly non-social and only interacted when they happened to use the same areas simultaneously, randomizing the data in this fashion would likely yield the same amount of co-location events. The only assumption that this null model makes is that key resources which drive mobility do not fluctuate significantly on the scale of days. See (Spiegel et al., 2016) for an in-depth discussion of the null model.

Finally, we acknowledge that a stronger statistical approach would have been to produce many (thousands) of such shuffled datasets with this null model and report average effect sizes as well as *p*-values associated with each effect. For the current dataset this was not computationally feasible, since we inferred interactions by querying the distance between every pair in every time-step, which yields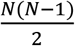. 2.1 · 10^6^ timesteps ≈ 1.3billion queries (or ~2 days of computing time using 56 2.60GHz processors) per shuffled dataset.

### Timescale Analysis to Identify Communities

We conducted a computational experiment where we measured how the number of communities changed when we incrementally split each monthly social network into multiple shorter aggregates. Specifically, we first aggregated all interactions in a four week window within a given month and weighted links by the number of interactions between two fish that exceeded the background level of interaction (number of interactions in the null data). We then broke this network into two two-week networks each mapping the interactions in their given window. We continue breaking up the networks into an increasing number of temporally shorter networks until the window size was one hour and the number of networks was 24 h/day · 28 days=672. In each iteration we ran the community detection algorithm Infomap (Rosvall, Axelsson, & Bergstrom, 2009) on the networks and recorded the number of communities that had three or more members. In the results, we report the average and standard error of the mean across slices for each aggregation level.

## Results

### Behavioral trends during day and night across the year

#### Behaviour was variable across the season, while strong shoaling was a daytime winter phenomenon

We observed high variation across most individual and social behaviors across the year. This variation was largely due to the fish shoaling in deep waters at the center of the lake during daytime in colder months. We saw differences in swimming speed (Jan. avg.: 1.03 m/s, June avg.: 0.51 m/s) depth (Jan. avg.: 5.15 m, June avg.: 1.48 m) and distance from shore (Jan. avg.: 103.38 m, June avg.: 40.22 m). Moreover, we found that indicators of social interactions in the colder months were elevated, such as time spent together (Jan. avg.: 12.22 minutes, June avg.: 9.32 minutes), and interaction probability (Jan. avg.: 0.154, June avg.: 0.042). In colder months there was also a great difference between social indicators during day and night, which we interpret as a strong signal that shoaling is indeed a daytime phenomenon in this population. Figure 2 contains a full summary of these results. In warmer months starting late March, as the lake stratified (Figure SI2), deep waters turned anoxic forcing the carp closer to the surface. We observed already in late February that interaction probability and duration of interaction during daytime decreased, while time between interactions increased overall. This means that shoaling ceased before deeper water layers turned anoxic in spring and over the summer. Shoaling in daytime re-appeared gradually during the fall and peaked again in December.

**FIG. 2.**
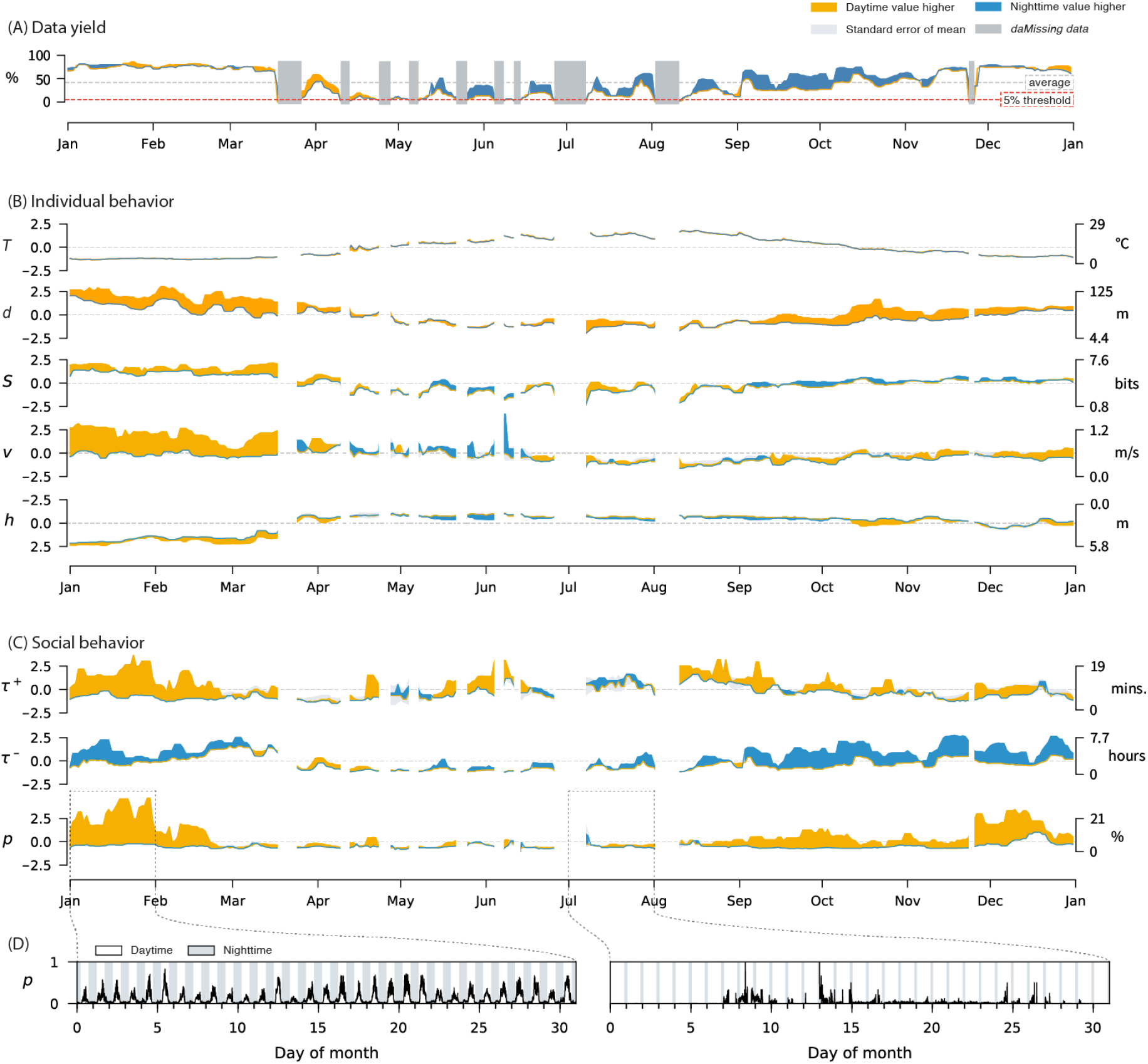
Seasons and daytime modulate individual and social behavior. (A) Percentage successful location measurements across the year. Using a 5% threshold we label the periods marked in grey as missing data. Notice how these periods are missing in panels (B) and (C). (B) Individual behavioral indicators computed from location data, including temperature, T, distance from shore, *d*, spatial entropy, *S*, velocity, *v*, and depth, *h*. All values in panels (B) and (C) are population averages. Deviations are represented as the average standard error of the mean of day and night, which rarely exceeds the absolute difference between day and night. The left set of axes are standardized values representing standard deviations from the mean, and the right give the actual values. (C) Social behavioral indicators measured at the level of pairs. We measure social activity in terms of the typical interaction duration, +, time between interactions, -, and interaction probability, *p*. (D) Example of how *p* varies rhythmically throughout January and July.

#### Behavioral differences between day and night were seasonal

Across all the behavioral and social indicators we studied, we only observed high variation over daytime during the winter, but not in the other seasons. Comparing the time-series of *p* (interaction probability) for January with July, it is clear that social interactions were a periodic function of daytime only in the winter, whereas they were far more sporadic in the summer (Fig. 2D). A notable exception, though not directly an indicator of behavior, was data yield. Here, we observed the opposite: in winter, roughly the same amount of location measurements were successful across the solar cycle, whereas in summer, the fish were significantly easier to detect (higher yield) at night (Fig. 2A). This is surprising because we simultaneously observed the carp to swim closer to the shore during the night (lowered *d*, Fig. 2B), where we would expect more signal attenuation due to reed growth, suggesting that the carp actively swam in denser vegetation during daytime in the summer, moving to the sublittoral areas during night where detectability increased.

#### In summer, interactions are less frequent but more persistent

In warmer months, when the deep zones of the lake become anoxic and the food was concentrated in the littoral zone, the carp had little to gain foraging-wise from swimming at the lake centre or in the deep water in the lake centre. Instead, they resided in the shallow waters alongside the shore where they could seek protection from predators among the food-rich reeds without investing much energy in mobility (lowered *d*, *S*, and *v*, Fig. 2B). This introduced a technical inconvenience as reed attenuated acoustic signals, which caused the data yield to drop substantially during summer (Fig. 2A). Despite this, we can report the somewhat surprising result that during the summer when carp associated strongly with the vegetated littoral zone, they interacted less often (lowered *p*, Fig. 2C) but also spend less time apart between interactions (lowered *τ*^-^, Fig. 2C). Additionally, *τ*^+^, which we should expect to drop across the summer due signal attenuation, appeared, although noisy, stable throughout the year. This suggests that inside the reed, fish were interacting in small persistent groups.

### Social attraction or co-use of suitable habitats?

#### Social attraction was a key driver of social interactions

We found that across the year, there were one to six times more interactions in the real data than in the shuffled data (Fig. 3B). This is strong evidence that social attraction was, in most months of the year, a key driving mechanism for proximity interaction of carp. The effect sizes were greatest at the start of the year in winter and decreased towards autumn. The overall number of interactions in the real data dropped as well (Fig. 3A, note that y-axis is log-scaled). We also observed that effect sizes were larger during daytime than at night (Fig. 3B). This indicates that nightly mobility (or *stationarity*) took place in the same locations over at least a week and was less driven by where other fish spend their nights.

**FIG. 3.**
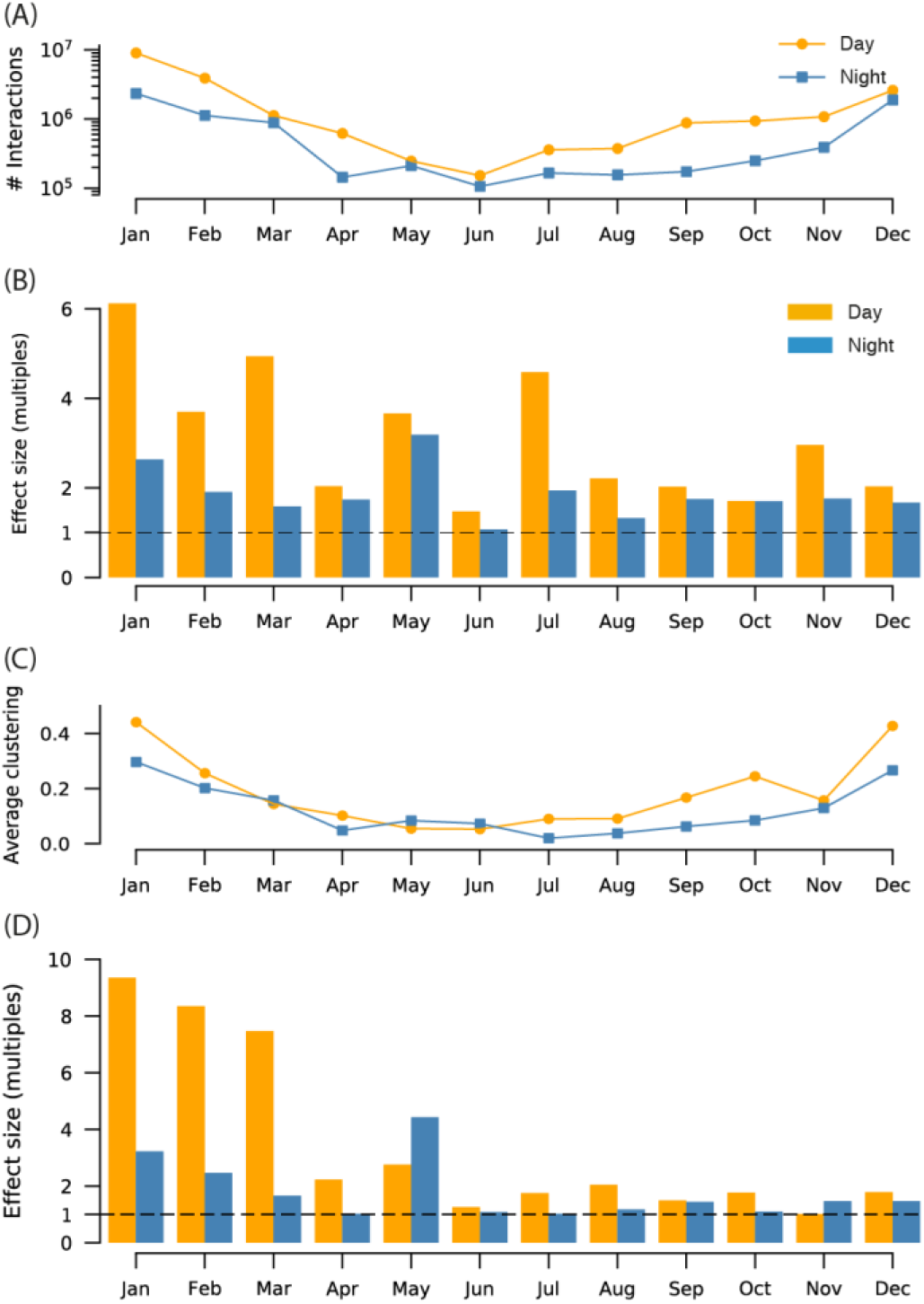
Effect sizes compared to shuffled data. (A) Number of interactions observed during day and night in the real data over the year. Note the y-axis is log-scaled. There are many times more interactions during daytime, and a decrease in total number of interactions observed over the year. (B) The effect size of number of interactions in the real data as compared to the shuffled data (count of real divided by count of shuffled). The dashed line signifies a one-to-one ratio and therefore no effect. (C) Average network clustering coefficient. (D) Effect size of clustering.

#### Elevated local clustering revealed tight knit community structure

We found that clustering was higher during daytime than at nighttime in most months (Fig. 3). Clustering varied over the year, where we observed a decline in clustering from winter through summer and then an increase from autumn to winter, where by December, clustering had returned to the same level as in January, in spite of the number of interactions being many times smaller. Nighttime clustering also varied periodically in synchrony with the seasons. We measured the effect size of clustering by comparing to our null model, and observed that the effect sizes decreased over the year, to the point where we could not confidently state that any of the observed local clustering was due to social attraction. At the same time, however, the local clustering was just as high at the end of the year as it was in the beginning, leading us to reason that over the year the fish grew more habitual, visiting the same few locations every day at regular times.

### Network structure at varying time-scales

#### Social memory varied over the year

If the carp population had no collective social memory they would not form clusters, but instead mix randomly, and any emerging communities in the interaction network would be due to simultaneous space use. We studied the interaction networks that emerged when aggregating over time-windows of different duration (subtracting the corresponding random network produced by the null model), and measured the number of communities with three or more members as a proxy for the use of social memory in interactions (Fig. 4). We reason that when it is possible to aggregate over a long time window and still obtain a number of communities, there is a high degree of social memory in the population. Furthermore, the longest aggregation that does not wash out communities is a good estimate of the time-scale of social memory. We found that social memory was significantly more time-persistent in the summer where, for example, the number of detected communities at night peeked at the two week aggregation for months May and August. Reversely, in winter months like January and December, the network mixed on very short time-scales. This is not surprising, as we know this mixing happened due to shoaling, however, since the fish only shoaled during daytime we were surprised to find that that this mixing to a large extent persisted into the night.

**FIG. 4.**
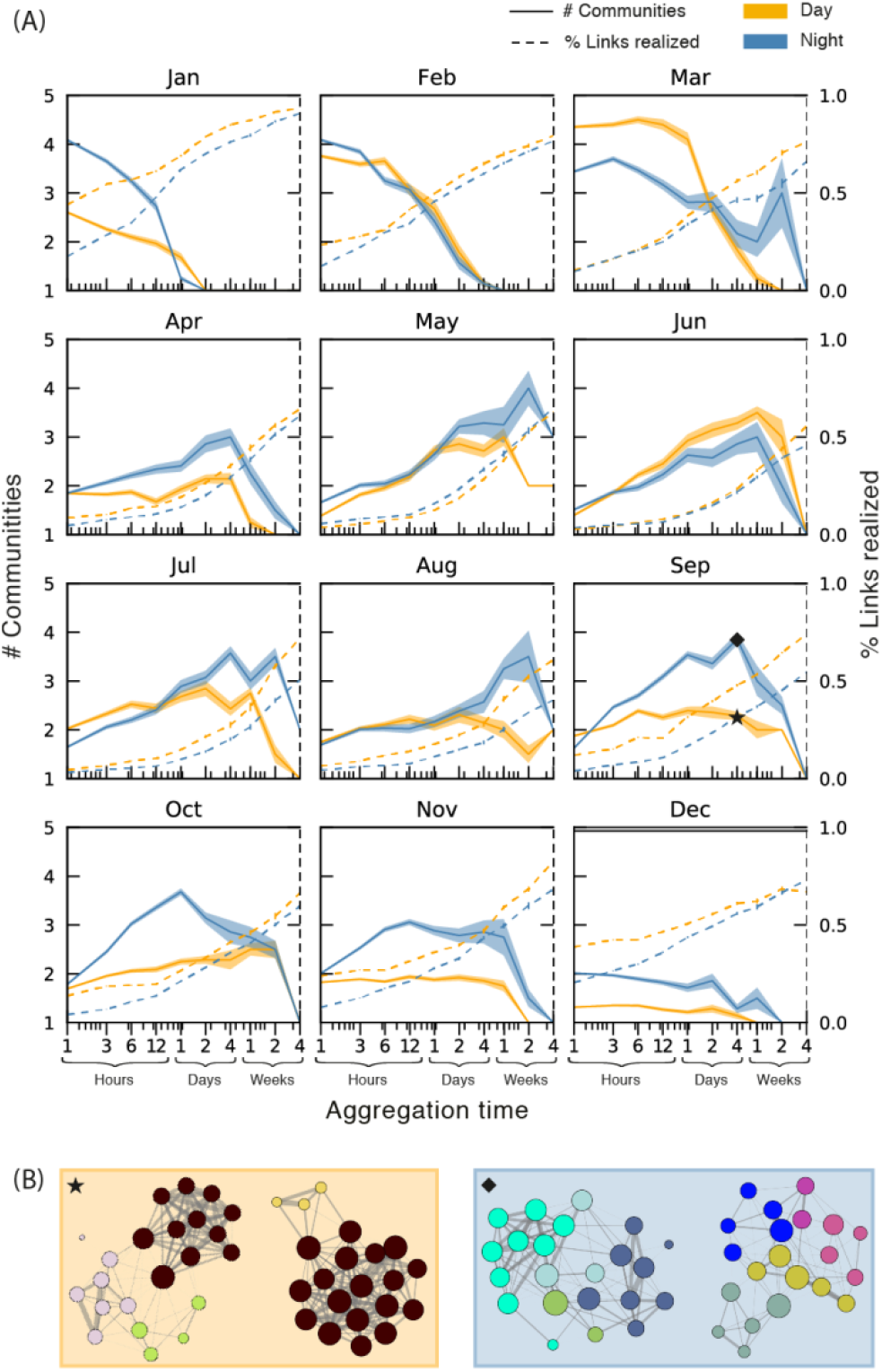
Number of communities at different levels of aggregation. (A) Number of communities (3 or more members) and percentage of realized links as a function of aggregation time for each month. Each has its own y-axis. Shaded regions surrounding the solid lines are standard errors. The x-axis is log-scaled. (B) Four example networks, two for day (left) and two for night (right), with communities detected (coloured separately) by the Infomap algorithm [18]. Each is an aggregate over four days in September, the references marked with a star and a diamond on the corresponding curves in (A).

**FIG. 5.**
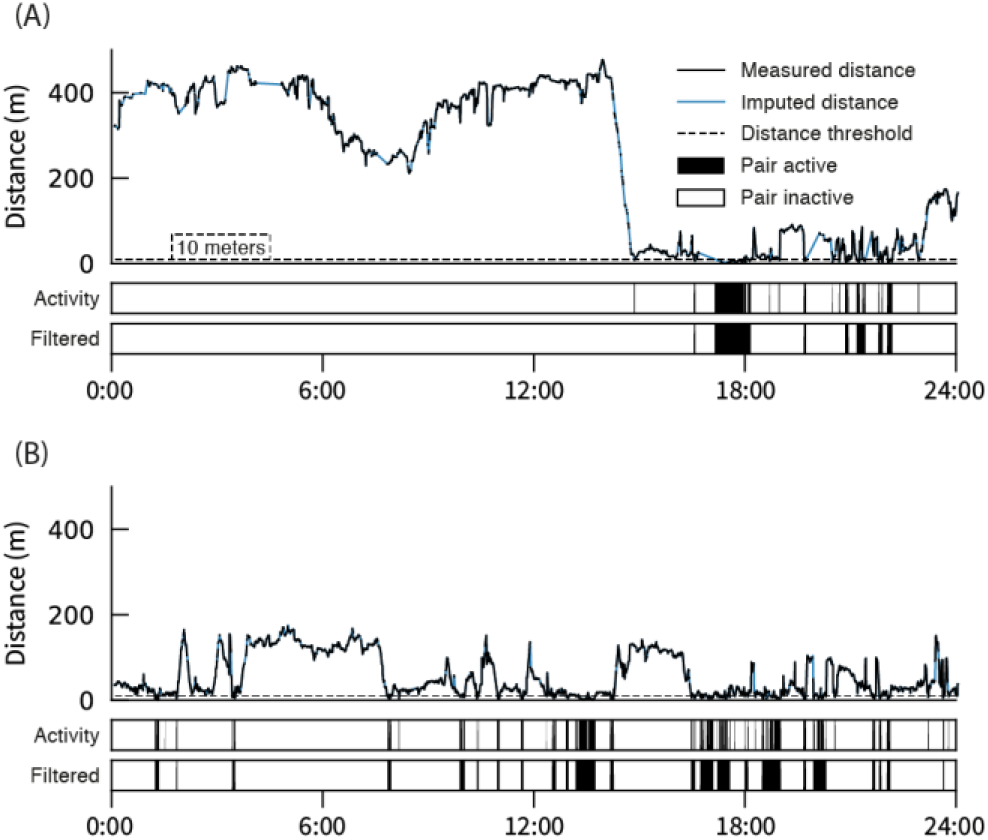
Inferring proximity network with distance threshold. (A) and (B) show how distance between two different pairs of fish vary throughout the same day. The dashed line at *Distance* = 10 m is the threshold we enforce to get the activity series shown below, which we filter to produce the bottom series.

#### Night communities were numerous and had longer time-scales

In nine out of 12 months, the number of communities during the night was higher than in daytime. This is not surprising because the fish interacted much more during the day, which caused mixing between existing groups on shorter time-scales. Curiously, however, the measured time-scales (aggregation time where number of communities peaks and then drops) of nightly communities were in many cases longer than those of day communities.

## Discussion

We observed that the social behaviour of a population of carp was highly dynamic across daily and seasonal scales, where the clustering and social memory was oscillating based on interactions between diurnal and seasonal rhythms. Thus, inferring social network information from time snapshots of data in selected weeks or months can lead to biased conclusions. Our analysis also showed that the groups of carp were indeed aggregating because of social attraction and the carp were likely to spend time in characteristic groups with the same individuals for up to two weeks particularly during the nighttime in the summer, indicative of pronounced social memory. We finally observed higher clustering during the daytime than the nighttime, and surprisingly large shoals of actively moving carp in the wintertime during daytime, which as discussed below is likely a response to environmental conditions in the lake and the origin of the fish. Methodologically, our analysis can serve as a template for future investigations into the causes and consequences of social behaviour in the wild as more high-resolution movement data in the wild begins to emerge. Our findings can also help inform both sampling design (Bajer & Sorensen, 2012; Muška et al., 2018) or removal strategies (Bajer et al., 2011; Carl, Weber & Brown, 2016) for common carp in the wild.

We found highly variable social behaviour of our carp population in the wild over the course of a full year. The carp aggregated in somewhat deeper zones in wintertime in agreement with previous natural history knowledge of the species (Armstrong et al., 2016; Bajer et al., 2011; Bauer & Schlott, 2004; Gusar, 1989; Johnsen & Hasler, 1977; Penne & Pierce, 2008; Taylor et al., 2012), however, these zones did not encompass the deepest points of the lake. Previous work in other systems with low resolution telemetry has reported carp were largely sedentary during the wintertime with only localized activity (Bauer & Schlott, 2004; Johnsen & Hasler, 1977; Jurajda et al., 2016; Penne & Pierce, 2008), in stark contrast to the high levels of daytime activity we observed during winter in Kleiner Döllnsee. Elevated levels of overwintering activity of carp are usually attributed to stressors, such as low oxygen (Bauer & Schlott, 2004), movement of humans on the ice surface (Johnsen & Hasler, 1977), or predators (Adámek, Sukop, Moreno Rendón, & Kouřil, 2003). In our study, hypoxic conditions was an unlikely stressor as Kleiner Döllnsee is well oxygenated in the wintertime. Moreover, the fish were not manually tracked eliminating human on-ice activity as a possible disturbance. Therefore, reasons for the surprisingly active shoaling behaviour are likely related to “hypothetical” predator avoidance behavior shown by the introduced carp in their new, unfamiliar environment particularly during daytime in response to visual predators. Shoaling is a common response to predation risk (Pitcher, 1986) as it offers increased predator detection probability (Godin, Classon, & Abrahams, 1988) and dilution of risk (Queiroz & Magurran, 2005), and shoaling can confuse predators (Krakauer, 1995). The carp we tracked were still likely responsive to predation risk despite the fact that they were large enough to escape the threat of predation from most predators in the lake (Gaeta et al., 2018). Possible predators include European catfish (Carol, Benejam, Benito, & García-Berthou, 2009), great cormorants (*Phalacrocorax carbo*) (Adámek, Kucerova & Roche, 1999) or mustelids such as the European otter (*Lutra lutra*) (Adámek et al., 2003; Britton, Pegg, Shepherd, & Toms, 2006). Indeed, carp in a reservoir were observed to spend their nights in deep hypoxic waters, which was also speculated to be a response to a population of wels catfish present in the reservoir, which hunt nocturnally (Benito et al., 2015). We found that the carp reduced their shoaling behaviour in the second winter in our study, after they were more familiar with the (rather low predation) risks of the novel environment, which lends further support to the idea that daytime winter shoaling in the first winter after introduction to their new environment was likely a predator-avoidance response driven by unfamiliarity with possible predation threats during daytime in winter. We expect that carp should behave cautiously in a novel lake environment, despite a lack of strong predation risk due to their larger size (Lorenzen, 2000). The anti-predator behaviour could become fixed from early life experience or from the evolutionary past (Blumstein, 2006; Magurran, 1990; Swaney, Cabrera-Álvarez, & Reader, 2015). When compared to behaviour in a controlled laboratory environment conducted in large tanks, carp have been observed to behave highly cautiously after introduction to a predator free semi-natural pond environment of comparable size, where the fish reduced their visits to open feeding sites (Klefoth et al., 2012). In the case of the experiment by Klefoth et al. (2012) water from a nearby lake was flowing through the semi-natural ponds, and therefore the carp were likely exposed to chemical cues signalling that predators could be present. Hence, it is reasonable that the fish tracked in our experiment, which were recently introduced to the lake, could be behaving cautiously, despite a low actual predation risk. After learning about the true predation threat in the lake was likely low (in fact no otters were recently seen in the lake and cormorant predation is low too), the carp likely behaved less cautiously in the second winter and reduced their daytime social behavior substantially.

The carp were more likely to shoal in the daytime than the nighttime throughout most of the year. As the daytime clustering corresponded with offshore movements, into riskier habitat outside plant refuges, the daytime clustering may partially be explained as a response to visual predators such as cormorants (White et al., 2008) or pike (Eklöv, 1992) that actively hunt during daytime in the study lake (Kobler et al., 2008). Diurnal migrations, in particular diurnal vertical migrations, are also a well known behavioural response displayed by many smaller bodied fish species, macroinvertebrates and zooplankton, where individuals balance bioenergetic efficiency, foraging opportunities and predation risk, by sheltering in deeper water and foraging in shallower waters when predators are less active (Mehner, 2012). Some cyprinids have also been observed to migrate horizontally on a daily scale (Kubečka, 1993; Nakayama et al., 2018), as we observed the carp doing in our study. Similar to the carp, common bream (Schulz & Berg, 1987) and freshwater drum, *Aplodinotus grunniens* (Rypel & Mitchell, 2007), have been observed to move from littoral habitats during the night to pelagic habitats during the daytime. Further a whole fish assemblage in a Czech reservoir aggregated in the pelagic during the daytime and spread out in the littoral during the nighttime (Muška et al., 2013, 2018). Other small cyprinid species have been observed to migrate horizontally in the opposite direction to our tracked carp, sheltering in the littoral during the daytime and foraging in the pelagic at nighttime (Haertel & Eckmann, 2002; Nakayama et al., 2018). The ultimate mechanism for diurnal horizontal migrations and in which direction they occur in these species is not known, but it is expected to also relate to a tradeoff between resource availability and predation risk (Rypel & Mitchell, 2007; Schulz & Berg, 1987; Shoup, Boswell, & Wahl, 2014). Hence, in the summertime the carp may be foraging in the littoral habitat during the night-time, avoiding predation risk from the pelagic catfish (Benito et al., 2015; Carol, Zamora, & García-Berthou, 2007) and foraging in more pelagic habitat during the daytime, while forming shoals to reduce perceived predation risk from pike or otters.

We found longer-lasting and smaller groups of fish during the summertime, in particular during the nighttime. Our comparisons to a null model of behaviour (Spiegel et al., 2016) indicated that these clusters were not driven by attraction to similar locations at similar times, but were truly a result of attraction to the individual carp. We found that the carp during the summertime had a pronounced social memory, showing preferences to interact with certain individuals for up to two weeks. Importantly, two weeks is longer than it took the twelve days for guppies, *Poecilia reticulata*, to learn and retain the identity of conspecifics (Griffiths & Magurran, 1997); hence it is likely that the carp were able to remember the identities of their conspecifics. Furthermore, although the carp spent their days mixing in larger groups, they tended to spend the nights together in smaller characteristic groups. Carp anglers have long noted that carp have “friends”, as many have observed that after catching one specific individual carp a second individual is predictably captured (Hearn, 2000). In juvenile fish preferred interactions have can be based on kinship (Piyapong et al., 2011); however, kinship is typically not the case in adult fish (Croft et al., 2012; Russell, Kelley, Graves, & Magurran, 2004), and therefore it is likely that the groups we observed were not based on kinship. In the summer months, Kleiner Döllnsee is productive and resource-rich; however, the spatiotemporal distribution of food is patchy across the lake. In such patchy environment, carp should benefit from information sharing (Monk et al., 2018) to find resources faster. Indeed, when feed bags were introduced into a lake the whole population of carp were able to find a feed bag within four nights, much faster than possible by individual private searching (Bajer et al., 2010), suggesting social learning and other forms of communication (e.g., chemical communication through excretion; Brönmark, & Hansson, 2000). Therefore, we suggest that the small groups of carp we found in the summer may provide them with valuable information regarding resource distribution in the lake. Carp are also quick to learn from trained demonstrators (Karplus et al., 2007; Klefoth et al., 2012; Zion et al., 2007), and can retain socially learned information for at least one year as, population-wide catch rates were found to decline for an entire year after angling a pond despite most individuals in the population never having been captured (Beukema, 1970).

Recognizing specific individuals by social memory may provide additional foraging benefits to the carp in our study. Familiarity in general is known to provide fitness benefits (Seppä, Laurila, Peuhkuri, Piironen, & Lower, 2011), in particular via the increased foraging success through directed social learning as fish may learn better from familiar individuals (Swaney et al., 2015). Hence, knowing and following the most productive foragers in the population should be beneficial, especially in the context of producer scrounger dynamics, where some individuals may be generally better at finding new food sources through private information, while other individuals tend to follow those individuals to find food (Caraco & Giraldeau, 1991). As well, carp are known to show consistent inter-individual differences in foraging rate (Pollux, 2017) and individual variation in diet (Mehner et al., 2018) and taste preferences (Kasumyan, 2000). Consequently, certain individual fish may have better information regarding certain food sources within the patchy resource distribution in a natural lake, providing fitness benefits to sociality during resource-rich environmental conditions in the warmer periods of the year.

## Conclusion

We found that carp adjust their social behaviour following several timescales of oscillation, where yearly and daily oscillations very likely respond to variation in perceived predation risk and resource availability, moderated by the benefits of social interactions. Despite a low realized threat of predation in our study lake given the large size of the tagged carp, the fish displayed cautious behaviour after introduction into a novel environment, particularly during the day and in winter by revealing strong tendencies of shoaling. Further, the carp displayed pronounced social interactions based on social attraction with the community organization being non-random and based on a social memory. These findings strongly indicated that carp are able to recognize one another and to take advantage of familiarity during productive phases in warmer months of the year. To our knowledge, our work is among the first year long high-resolution analyses of animal social networks in the wild. Our analysis may serve as a methodological template for future analyses into the rhythm of relationships in other species and taxa.

## Acknowledgements

This work was supported by a Leibniz Community (“B-types”, SAW-2013-IGB-2) grant received by RA. This work was also supported by Strategic Grants by Princeton University and Humboldt-Universität zu Berlin on “Princeton-Humboldt Centre for the Reality Mining of Animal Systems,” the ‘Cooperation and Collective Cognition Network’. We would like to thank Andreas Mühlbradt, Alexander Türck, Jan Hallermann, Jacob Weinrautner, Jonathan Nickl, Bernard Chéret and many other technicians and students for help in the field and processing the data. The experiments were approved through animal care permit (2347-21-2014) granted by the Ministry of Environment, Health and Consumer Protection Brandenburg, according to the German Animal Protection Act.

