## Supplemental Information for "Rhythm of relationships in a social fish over the course of a full year in the wild"

### Short Title: Dynamics of Carp Social Networks

**Ulf Aslak[1*], Christopher T. Monk[2*], Dirk Brockmann[3,4], Robert Arlinghaus[2,5,6]**

*[1]Centre for Social Data Science, University of Copenhagen, DK-1353 København K and DTU Compute, Technical University of Denmark, DK-2800 Kgs. Lyngby. [2]Department of Biology and Ecology of Fishes, Leibniz-Institute of Freshwater Ecology and Inland Fisheries, Müggelseedamm 310, 12587 Berlin, Germany. [3]Robert Koch-Institute, Nordufer 20, D-13353 Berlin, Germany. [4]Institute for Theoretical Biology and Integrative Research Institute for the Life Sciences, Humboldt Universität zu Berlin, Berlin, Germany. [5]Faculty of Life Sciences and Integrative Research Institute for the Transformation of Human-Environmental Systems, Humboldt-Universität zu Berlin, Invalidenstrasse 42, 10115 Berlin, Germany.[6]Division of Integrative Fisheries Management, Department of Crop and Animal Sciences, Faculty of Life Science, Humboldt-Universität zu Berlin, Invalidenstrasse 42, 10115 Berlin, Germany.*

***** These authors contributed equally to this work

**Supplemental Information**

**Table SI1.** A summary of all acoustically tagged carp that generated data for our temporal networks, including size at tagging and data yield.

| ID | Total Length (mm) | Wet Mass (g) | Daily Detection | | | | |
|---|---|---|---|---|---|---|---|
| | | | Mean ± sd | Minimum | Maximum | Mean % | Maximum % |
| 59800 | 674 | 5596 | 2955 ± 2223 | 0 | 7574 | 7.10 | 43.83 |
| 60300 | 629 | 3903 | 2890 ± 2093 | 0 | 6715 | 6.72 | 38.85 |
| 61000 | 555 | 3170 | 2939 ± 2010 | 0 | 6791 | 7.00 | 39.29 |
| 61100 | 643 | 4127 | 1144 ± 1900 | 0 | 8484 | 6.62 | 49.09 |
| 61400 | 628 | 4028 | 3365 ± 2378 | 0 | 7912 | 9.47 | 45.78 |
| 61600 | 573 | 3323 | 3028 ± 2259 | 0 | 7247 | 7.52 | 41.93 |
| 62100 | 644 | 4779 | 2597 ± 2204 | 0 | 7270 | 5.02 | 42.07 |
| 62300 | 486 | 1840 | 1208 ± 2410 | 0 | 8163 | 6.99 | 47.23 |
| 62500 | 653 | 4429 | 1018 ± 1923 | 0 | 6979 | 5.88 | 40.38 |
| 62700 | 688 | 5159 | 1454 ± 2202 | 0 | 8238 | 8.41 | 47.67 |
| 62900 | 545 | 3440 | 1643 ± 2142 | 0 | 7537 | 9.50 | 43.61 |
| 63200 | 455 | 1850 | 2435 ± 2180 | 0 | 7051 | 4.08 | 40.80 |
| 63500 | 474 | 1530 | 1292 ± 2306 | 0 | 6862 | 7.47 | 39.71 |
| 63600 | 430 | 1171 | 3131 ± 2298 | 0 | 7204 | 8.12 | 41.68 |
| 63700 | 492 | 1783 | 3023 ± 2438 | 0 | 8720 | 7.49 | 50.46 |
| 63900 | 505 | 2130 | 1104 ± 2028 | 0 | 7012 | 6.39 | 40.57 |
| 64200 | 463 | 2398 | 1232 ± 2107 | 0 | 6629 | 7.13 | 38.36 |
| 64800 | 510 | 1760 | 3141 ± 2468 | 0 | 8325 | 8.17 | 48.17 |
| 64900 | 623 | 4083 | 2418 ± 2004 | 0 | 6527 | 3.99 | 37.77 |
| 65000 | 470 | 1700 | 1288 ± 2382 | 0 | 7769 | 7.45 | 44.95 |
| 65500 | 588 | 3367 | 2871 ± 2261 | 0 | 8395 | 6.61 | 48.58 |
| 65800 | 707 | 5872 | 1244 ± 2290 | 0 | 7258 | 7.19 | 42.00 |
| 66200 | 458 | 2230 | 3338 ± 2411 | 0 | 8009 | 9.31 | 46.34 |
| 67200 | 550 | 2633 | 3168 ± 2419 | 0 | 7441 | 8.33 | 43.06 |
| 67700 | 473 | 2451 | 3184 ± 2119 | 0 | 9584 | 8.42 | 55.46 |
| 67800 | 563 | 2915 | 2898 ± 2303 | 0 | 6824 | 6.77 | 39.49 |
| 68100 | 469 | 1794 | 1865 ± 1922 | 0 | 9166 | 0.79 | 53.04 |
| 68200 | 522 | 2439 | 1474 ± 2600 | 0 | 12855 | 8.52 | 74.39 |
| 69400 | 434 | 1354 | 2806 ± 2185 | 0 | 7211 | 6.23 | 41.73 |
| 69500 | 540 | 3227 | 3197 ± 2344 | 0 | 8108 | 8.49 | 46.92 |
| 69600 | 458 | 1647 | 3306 ± 2363 | 0 | 7585 | 9.13 | 43.89 |
| 69800 | 519 | 2041 | 2828 ± 2231 | 0 | 7812 | 6.36 | 45.20 |
| 70200 | 508 | 2213 | 1632 ± 2569 | 0 | 8051 | 9.44 | 46.59 |
| 70400 | 550 | 2810 | 1334 ± 2347 | 0 | 6822 | 7.72 | 39.47 |
| 70700 | 527 | 1791 | 653 ± 1198 | 0 | 6551 | 3.77 | 37.91 |
| 74000 | 437 | 1250 | 3292 ± 2419 | 0 | 7830 | 9.04 | 45.31 |

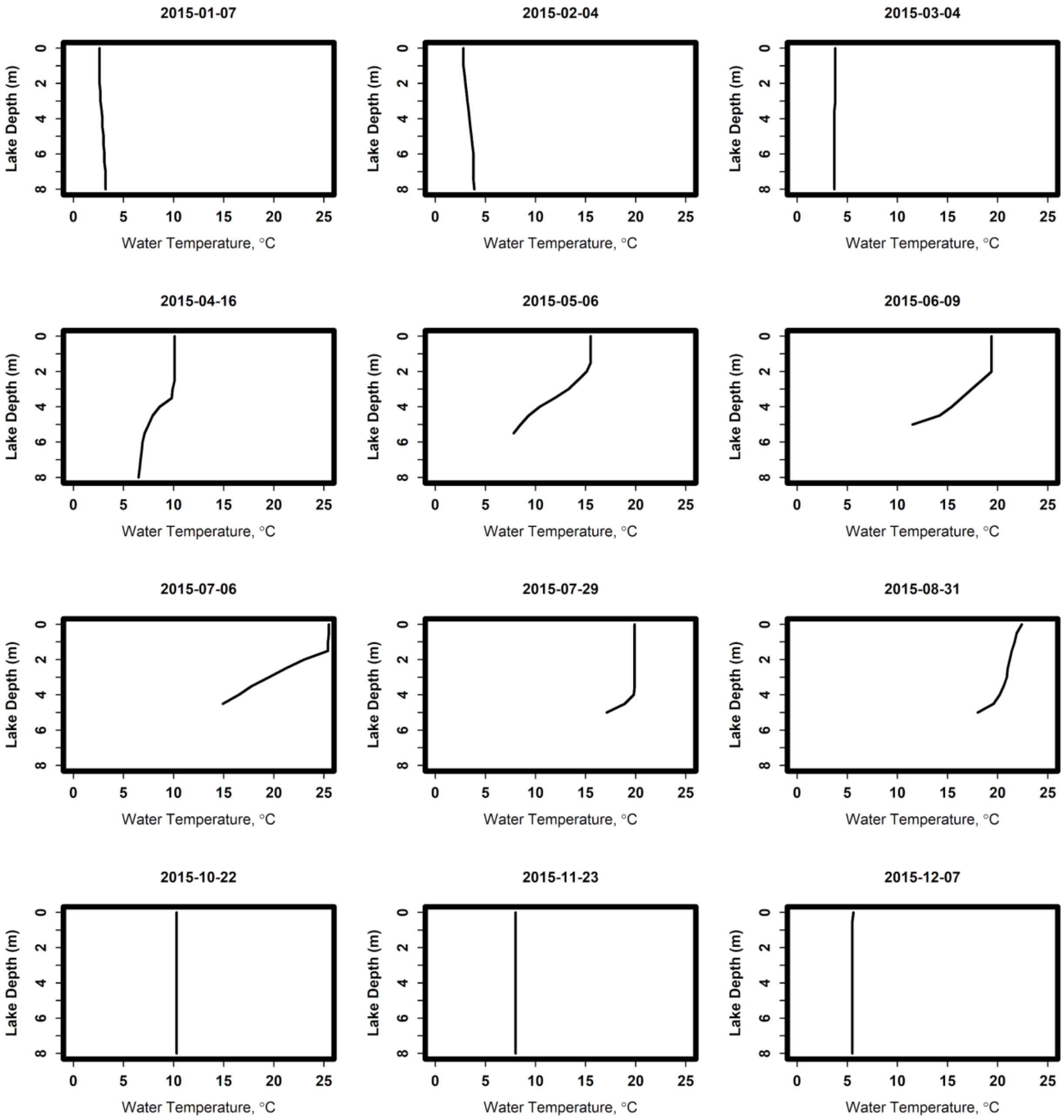


**FIG SI1** The temperature profile of the study lake, Kleiner Döllnsee for throughout the tracking period. The lake is stratified from May until October.

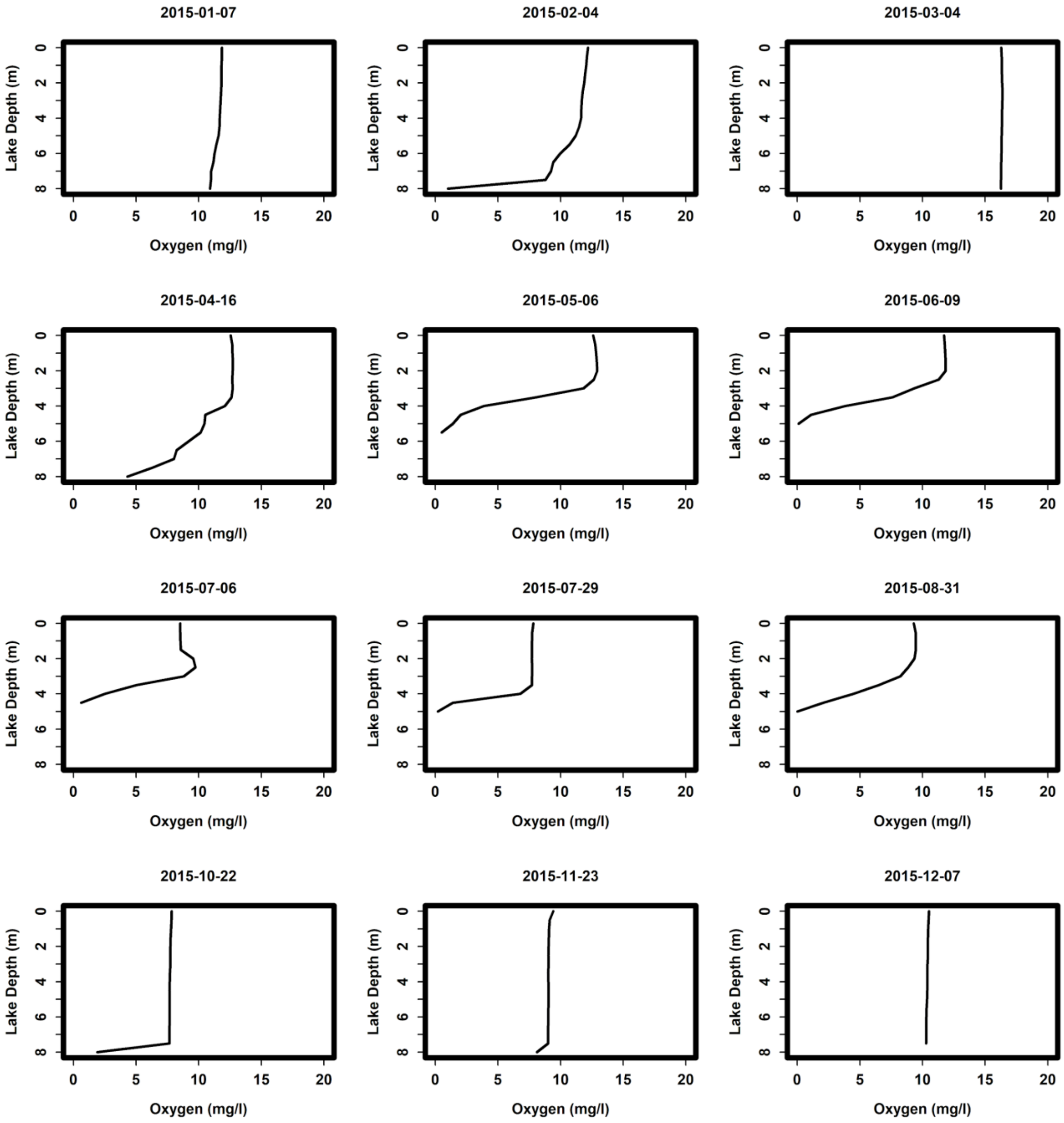


**FIG SI2** The oxygen profile of the study lake, Kleiner Döllnsee for throughout the tracking period. The lake is stratified from May until October, with an anoxic zone below approximately 4 m.
